# Structure of influenza D virus NS1 reveals a novel Zn-binding site and ssRNA-binding activity

**DOI:** 10.64898/2026.10.29.760789

**Authors:** Kai Ye, Minhui Guan, Andrew Kim, Mitch Miller, Rulong Ma, Clark Hamor, Aiden Wang, Ziqi Yuan, Bac Le, James M. Briggs, Xiu-Feng Wan, Yizhi J. Tao

## Abstract

Influenza D virus (IDV) is an emerging pathogen associated with bovine respiratory disease and has potential for zoonotic transmission. Although nonstructural protein 1 (NS1) is a major virulence factor in influenza A and B viruses, the structure and function of IDV NS1 remain poorly defined. Here, we determined crystal structures of the IDV NS1 N-terminal RNA-binding domain (NTD) and C-terminal effector domain (CTD). The NTD retains the conserved homodimeric six-helix fold, whereas the CTD adopts a distinct architecture containing a novel Zn²⁺-binding site. IDV NS1 CTD preferentially binds single-stranded RNA, and mutagenesis identified lysine and arginine residues critical for this activity. Single-residue substitutions disrupting either the Zn²⁺-binding site or RNA-binding interface prevented recovery of infectious recombinant IDV. These findings reveal distinct structural and functional features of IDV NS1 and establish the importance of CTD integrity for viral replication. Conservation of key CTD residues in influenza C virus NS1 suggests shared structural features that distinguish influenza C and D virus NS1 proteins from their influenza A and B counterparts.

## Introduction

The nonstructural protein 1 (NS1) encoded by influenza viruses is a multifunctional virulence factor that plays a central role in modulating host innate immune responses. Through protein-protein and protein-RNA interactions, NS1 has been shown to suppress antiviral signaling pathways and promote viral fitness [1]. In influenza A virus (IAV), NS1 binds double-stranded RNA (dsRNA) to antagonize the OAS/RNase L pathway and inhibit RIG-I-mediated recognition of viral RNA [2,3], suppresses activation of protein kinase R (PKR) [4], modulates apoptosis through interactions with cellular factors including p53 [5], Fas ligand [6], and phosphoinositide 3-kinase (PI3K) [7], inhibits host pre-mRNA processing through cleavage and polyadenylation specificity factor 30 (CPSF30) [8], and blocking host mRNA export to shut down host immunity [9]. Despite its critical role in immune evasion and pathogenesis, NS1 is dispensable for influenza A and B virus replication under certain experimental conditions [10–12], indicating that its primary role is to promote viral fitness rather than to directly mediate the viral replication process.

Structural studies have established a conserved architecture for NS1 proteins from influenza A and B viruses. NS1 proteins typically consist of four structural regions: an N-terminal RNA-binding domain (RBD or NTD), a flexible linker region, a C-terminal effector domain (ED or CTD), and a disordered C-terminal tail (CTT). Crystal structures of full-length proteins [13,14] and individual domains from multiple viral strains reveal that the NTD forms a homodimeric six-helix bundle responsible for dsRNA recognition [15], whereas the CTD adopts a characteristic β-sheet–helix fold that mediates interactions with host proteins and nucleic acids [16]. Despite extensive structural and functional characterization of NS1 from influenza A and B viruses, the corresponding proteins from influenza C and D viruses remain poorly understood, leaving an important gap in our understanding of the evolution and functional diversification of influenza virus immune antagonists, particularly given the substantial sequence divergence of NS1 proteins across influenza virus genera.

Influenza D virus (IDV) is a recently recognized member of the family *Orthomyxoviridae* that was first isolated from swine and cattle in 2011 [17] and subsequently classified as a distinct influenza virus genus [18]. Although IDV has been detected in multiple animal species, including swine, sheep, goats, horses, camelids, and white-tailed deer [19–24], cattle are considered its primary reservoir and a major driver of viral transmission [25]. IDV is an important contributor to bovine respiratory disease complex (BRDC), one of the most economically significant diseases affecting the cattle industry worldwide [20,26]. While no human disease associated with IDV has been reported, serological studies have demonstrated exposure among individuals with occupational contact with livestock [27]. Moreover, IDV replicates efficiently in human respiratory cell lines, human airway epithelial cultures, and lung tissues while suppressing innate immune sensing, supporting its potential for zoonotic transmission [17,28]. Understanding the molecular mechanisms by which IDV interacts with and evades host antiviral responses is therefore important for both animal health and preparedness for potential zoonotic emergence.

In this study, we determined the three-dimensional structures of the IDV NS1 RNA-binding domain (NS1D-NTD) and effector domain (NS1D-CTD). Despite limited sequence conservation with their influenza A and B virus counterparts, NS1D-NTD adopts the conserved structural fold characteristic of influenza virus NS1 proteins. In contrast, NS1D-CTD contains previously unrecognized structural features, including a Zn²⁺-binding site and a basic surface that appear to be conserved in influenza C virus NS1. Functional analyses demonstrated that disruption of either the Zn²⁺-binding site or the single-stranded RNA (ssRNA)-binding interface prevents recovery of infectious virus in MDCK cells under both normal and immunity-suppressed conditions, establishing an essential role for these features in IDV replication. Together, our findings reveal distinct structural and functional features of influenza D virus NS1 that appear to be shared with influenza C virus, providing new insights into the evolution and functional diversification of NS1 proteins across the influenza virus family.

## Results

### Biochemical characterization of NS1D

NS1D carrying an N-terminal His₆-thioredoxin fusion tag was expressed in *Escherichia coli* (*E. coli*) as a soluble protein (Fig. 1A). Purification using a heparin affinity column was essential for removing copurifying bacterial RNA. Following TEV protease cleavage of the fusion tag, NS1D eluted as a single peak during size-exclusion chromatography with an apparent molecular mass of ∼55 kDa (Fig. 1C, D), consistent with a homodimer and in good agreement with the calculated molecular weight of 54.8 kDa.

**Figure 1.**
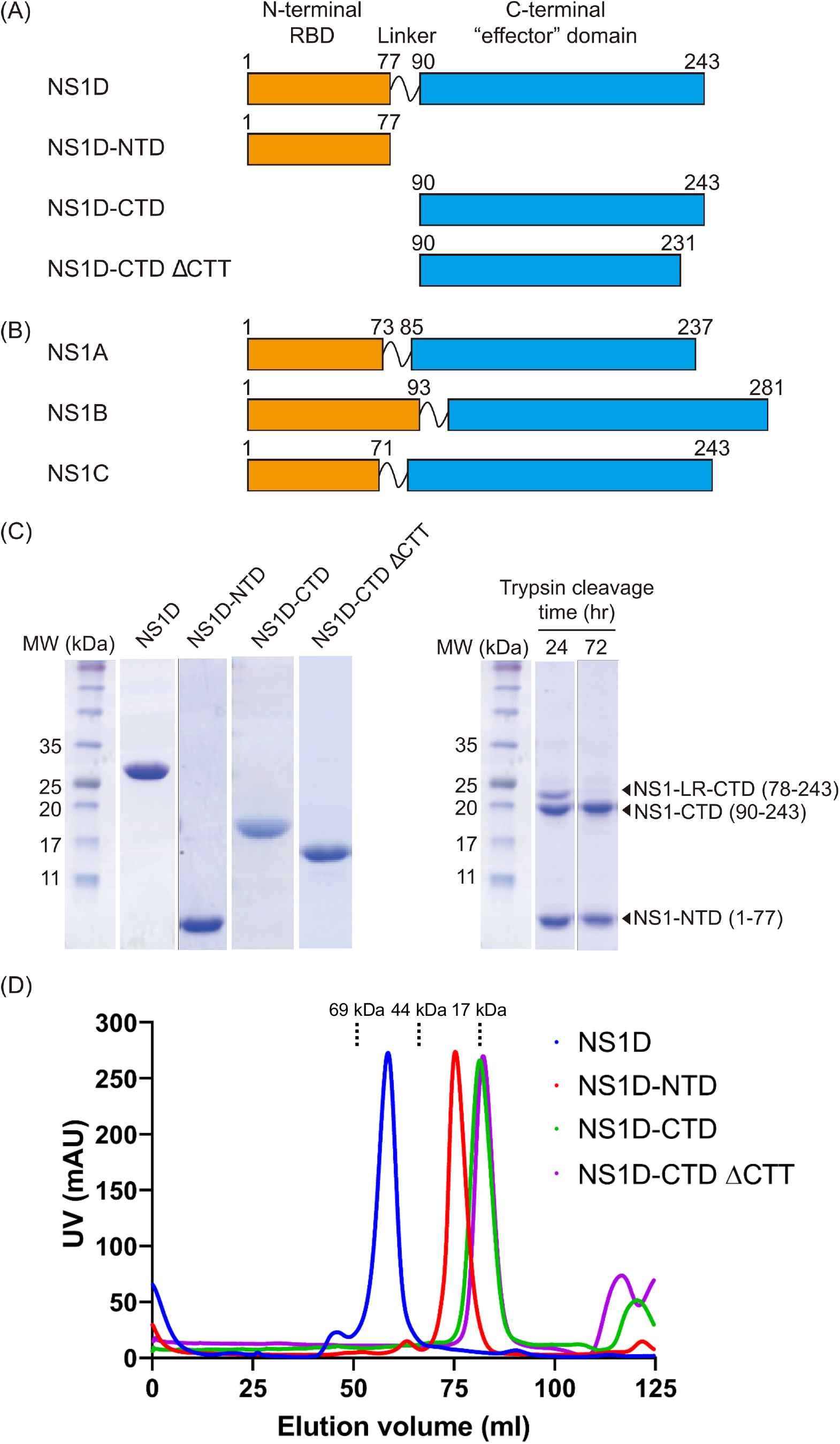
Recombinant NS1D constructs used in this study and their biochemical characterization. (A) Domain architecture of NS1D showing the boundaries of the N-terminal domain (NTD) and C-terminal domain (CTD). The amino acid ranges of the NS1D-NTD, NS1D-CTD, and NS1D-CTD ΔCTT constructs are indicated. (B) NS1 domain architectures from influenza A, B, and C viruses. Domain boundaries of NS1A and NS1B are based on available crystal structures, whereas those of NS1C are defined based on sequence comparison with NS1D. (C) SDS-PAGE analysis of purified NS1D proteins (left) and trypsin-digested samples (right). (D) Size-exclusion chromatography analysis of NS1D and its truncated constructs. Based on comparison with molecular weight standards, full-length NS1D and NS1D-NTD elute predominantly as homodimers, whereas NS1D-CTD and NS1D-CTD ΔCTT behave as monomers.

Despite its high purity and homogeneous dimeric state, full-length NS1D did not yield crystals in extensive crystallization screens. Because influenza virus NS1 proteins generally adopt a bilobed architecture (Figure 1B), we subjected recombinant full-length NS1D to limited trypsin digestion at a 1:100 molar ratio to define domain boundaries. Proteolysis generated two stable fragments after 72 h of digestion (Figure 1C, right panel). N-terminal sequencing of the cleavage products showed that the upper band corresponded to NS1D-CTD, beginning at residue 90 and extending to the C terminus, whereas the lower band corresponded to NS1D-NTD, comprising residues 1–77. A weaker higher-molecular-weight band observed after 24 h of digestion began at residue 78, suggesting that this intermediate fragment retained the flexible linker. NS1D-NTD and NS1D-CTD were subsequently subcloned, expressed, and purified from *E. coli*. Size-exclusion chromatography showed that NS1D-NTD and NS1D-CTD had apparent molecular weights of ∼17 kDa and ∼18 kDa, respectively, indicating that NS1D-NTD forms a dimer whereas NS1D-CTD remains monomeric in solution (Fig. 1D).

### Structure and dsRNA recognition by NS1D-NTD

NS1D-NTD (residues 1–77) crystallized in space group C2, and its structure was determined at 1.62 Å resolution by selenium single-wavelength anomalous dispersion (SAD) phasing [29] (Fig. 2, Fig. S1, Table 1). One NS1D-NTD dimer was present in each asymmetric unit. Each protomer consists of three interlocking α-helices (α1–α3) and a short C-terminal 3₁₀ helix (η1). Helices α1 (residues 5–26) and α2 (residues 30–52) are oriented nearly antiparallel to each other, whereas α3 (residues 57–68) is positioned approximately perpendicular to α1 (Fig. 2B).

**Figure 2.**
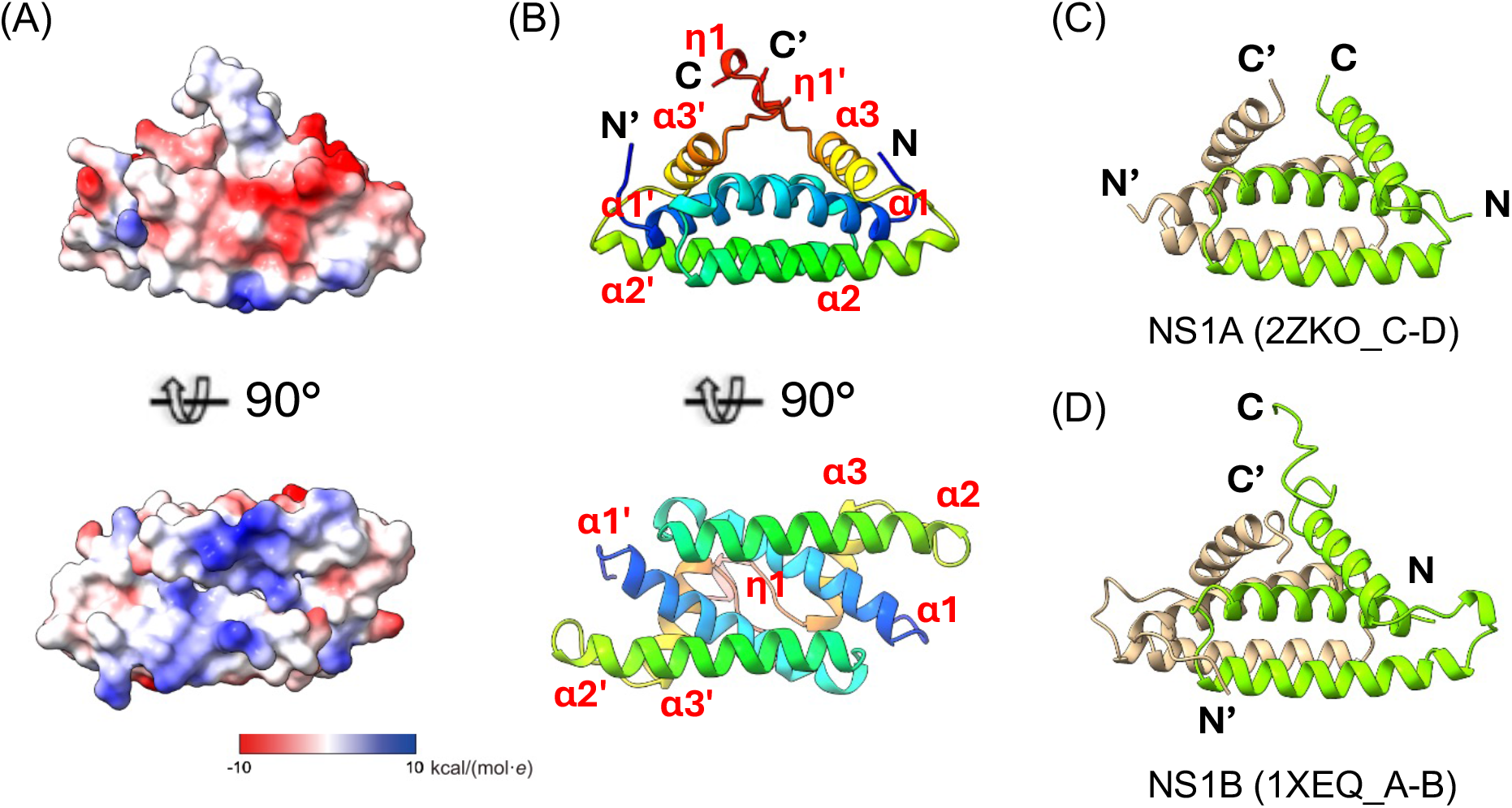
Crystal structure of NS1D-NTD. (A, B) Structure of the NS1D-NTD dimer. In the ribbon representation (B), both subunits are colored as a rainbow gradient from blue at the N-terminus to red at the C-terminus. In the electrostatic surface representation (A), positively and negatively charged regions are shown in blue and red, respectively. (C–D) Ribbon representations of NS1-NTD dimers from influenza A virus and influenza B virus. The two subunits are colored tan and green respectively.

**Table 1.** Data collection, phasing and refinement statistics

|  | NS1D-NTD (Se-SAD)<br>(residues 1-77) | NS1D-CTD<br>(residues 90-231) |
|---|---|---|
| PDB ID | 39II | 39IH |
| **Data collection** |  |  |
| Wavelength (Å) | 0.97856 | 1.19591 |
| Cell dimensions |  |  |
| *a*, *b*, *c* (Å) | 98.43, 58.24, 27.63 | 32.44, 60.49, 69.86 |
| α, β, γ (°) | 90.0, 94.7, 90.0 | 90.0, 90.0, 90.0 |
| Resolution (Å) | 50 – 1.62 (1.65-1.62) | 50-1.35 (1.44-1.40) |
| *R*<sub>merge</sub> (%) | 6.5 | 5.9 |
| *I* / σ*I* | 32.64 (5.77) | 17.61 (4.53) |
| Total reflections | 38752 (1793) | 165390 (6246) |
| Unique reflections | 19825 (1121) | 26916 (1511) |
| Completeness (%) | 98.4 (96.0) | 96.8 (75.9) |
| Redundancy | 1.9 (1.6) | 6.1 (4.1) |
| **Refinement** |  |  |
| Resolution (Å) | 28.51 – 1.62 (1.65 – 1.62) | 45.73 – 1.40 (1.44 – 1.40) |
| No. reflections | 19649 | 26916 |
| *R*<sub>work</sub> / *R*<sub>free</sub> (%) | 16.32/18.36 | 17.35/19.48 |
| No. atoms |  |  |
| Protein | 1233 | 1223 |
| Ligand/ion | 0 | 1 |
| Water | 137 | 162 |
| *B*-factors (Å<sup>2</sup>) |  |  |
| Protein | 15.86 | 22.11 |
| Ligand/ion | N/A | 13.53 |
| Water | 29.46 | 35.18 |
| R.m.s deviations |  |  |
| Bond lengths (Å) | 0.009 | 0.005 |
| Bond angles (°) | 1.224 | 0.925 |
| Ramachandran statistics |  |  |
| Favored (%) | 98.66 | 97.86 |
| Allowed (%) | 1.34 | 2.14 |
| Outlier (%) | 0.00 | 0.00 |

\*Values in parentheses are for highest-resolution shell.

The NS1D-NTD dimer adopts an overall triangular shape, with the two α2 helices forming the base of the triangle. The surface along this region is strongly positively charged, contributed in part by R13, K37, R39, and K40 (Figs. 2A & 3A).

**Figure 3.**
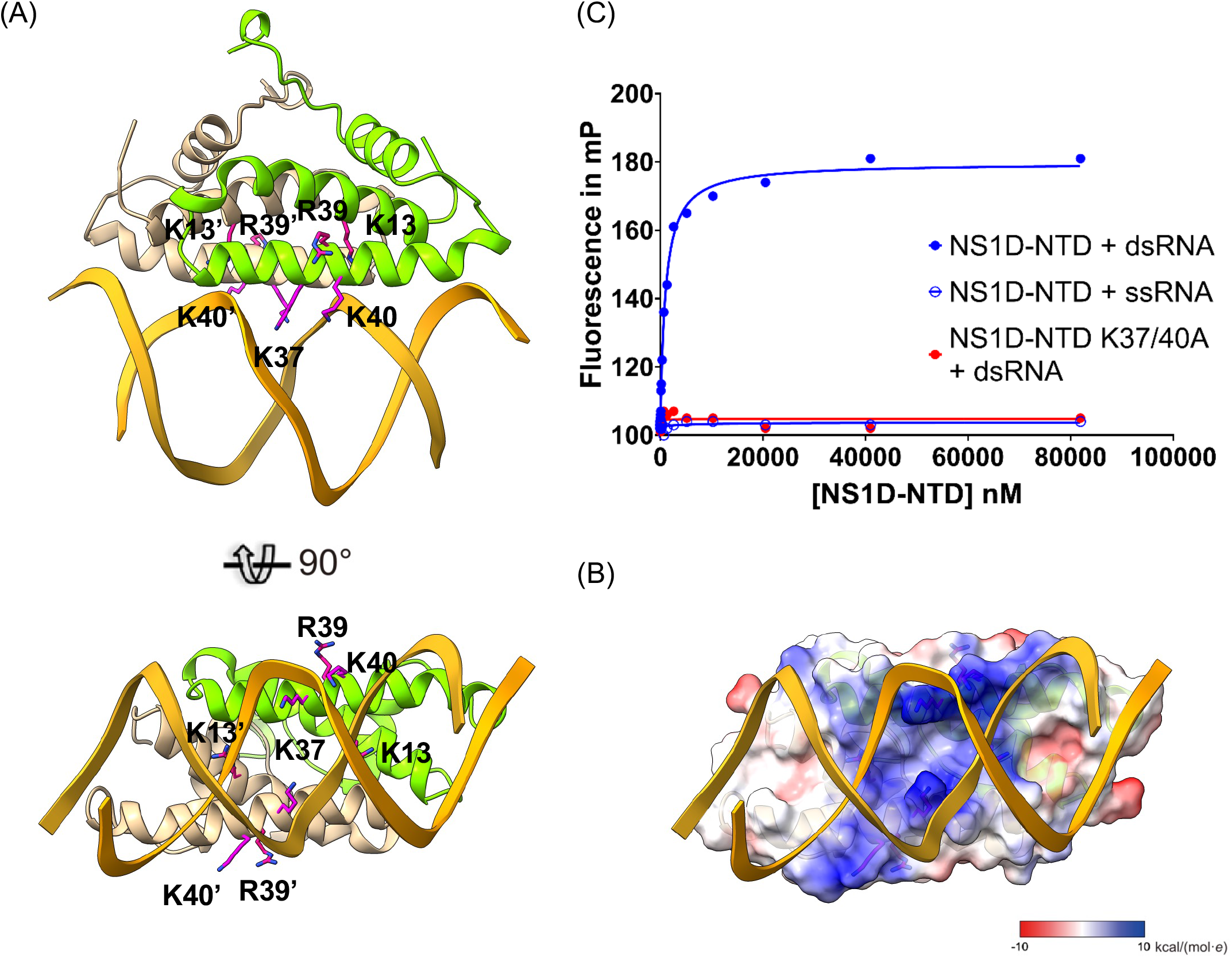
Molecular basis of dsRNA recognition by NS1D-NTD. (A) Structural model of the NS1D-NTD–dsRNA complex, with NS1D-NTD shown as a ribbon diagram (top) and as a surface representation (B). (C) Fluorescence polarization analysis of ssRNA and dsRNA binding by NS1D-NTD.

A Dali [30] search against the RCSB Protein Data Bank identified more than 40 NS1A and NS1B RBD structures with significant structural similarity. The top NS1A-RBD hit was PDB ID 2ZKO [31], with a Z score of 9.2, whereas the top NS1B-RBD hit was PDB ID 1XEQ [15], with a Z score of 9.6. Structural superposition of these NS1A- and NS1B-RBDs onto NS1D-NTD revealed a highly conserved homodimeric six-helix fold (Fig. 2C, D).

To model the interaction between NS1D and dsRNA, the NS1D-NTD structure was superimposed onto the H7N1 NS1A-RBD–dsRNA complex (PDB ID: 6ZLC) [32], which contains a 21-nt dsRNA molecule, and the resulting NS1D-NTD–dsRNA model was refined by energy minimization (Fig. 3). The model highlighted the importance of the positively charged surface in mediating dsRNA recognition (Fig. 3). In particular, the side chains of K37 and K40 project into the minor groove of the dsRNA and form hydrogen bonds with the RNA backbone, suggesting a central role in stabilizing the protein–RNA interaction.

A fluorescence polarization (FP) assay was used to assess the RNA-binding activity of NS1D-NTD using 21-nt synthetic RNA molecules labeled with Alexa Fluor 488. NS1D-NTD demonstrated moderate affinity for dsRNA (*K*_D_ = 3.0 ± 0.1 μM), but showed no detectable binding to ssRNA (Fig. 3B). The K37A/K40A double mutant showed no detectable binding to either dsRNA or ssRNA, suggesting that these residues play a critical role in mediating dsRNA binding by NS1D. This finding is consistent with previous studies showing that the equivalent residues in NS1A- and NS1B-RBDs are also crucial for dsRNA binding [33–35].

### Structure of NS1D-CTD reveals a unique Zinc-binding site

Purified NS1D-CTD did not yield crystallization hits. Because AlphaFold2 [36] predicted that the C-terminal tail (CTT; residues 232–243) is structurally disordered, we generated an NS1D-CTDΔCTT construct lacking the last 12 residues (Fig. 1). This truncated protein readily formed crystals, enabling structure determination by X-ray crystallography at 1.4 Å resolution, using molecular replacement with a structure model generated from a ColabFold prediction [37] (Table 1).

The NS1D-CTD structure comprises five β-strands and three α-helices, designated β1–β5 and α4–α6, respectively (Fig. 4 and Fig. S1). The five β-strands form a central twisted β-sheet against which the three α-helices pack, with α4 and α5 positioned on one side of the sheet and α6 on the opposite side. A Dali [30] search against the RCSB Protein Data Bank identified more than 40 homologous NS1A and NS1B CTD structures, with the top NS1A hit being PDB ID 3OA9 [38] (Z-score = 5.7) and the top NS1B hit being PDB ID 5DIL [16] (Z-score = 6.3).

**Figure 4.**
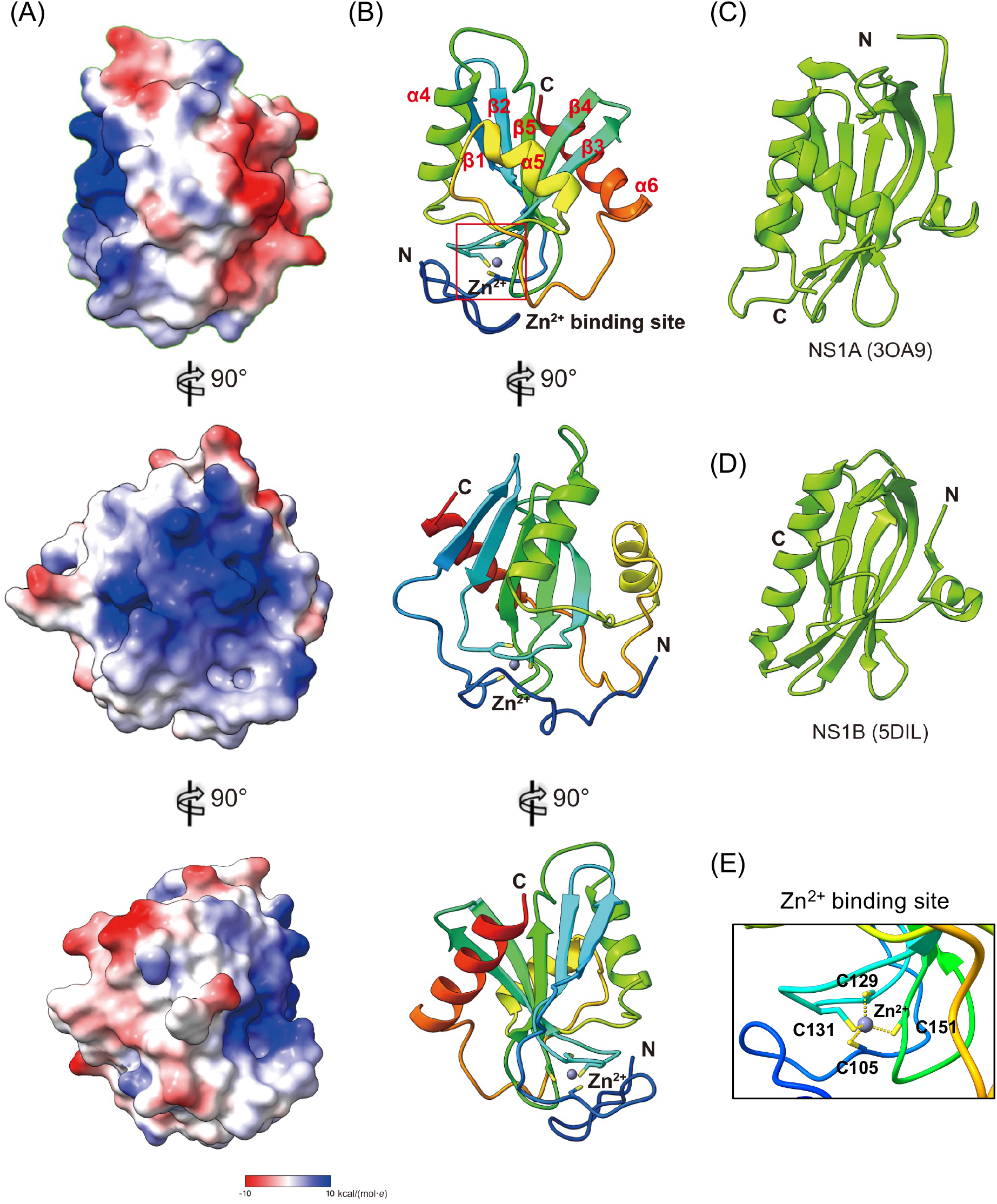
Crystal structure of NS1D-CTD. (A, B) Structure of the NS1D-CTD monomer. (A) Electrostatic surface representation in three orthogonal views, with positively and negatively charged regions shown in blue and red, respectively. (B) Ribbon representation colored in a rainbow gradient from blue at the N-terminus to red at the C-terminus. (C, D) Ribbon representations of the NS1A- and NS1B-CTD structures, respectively. (E) Enlarged view of the NS1D-CTD Cys₄:Zn cluster. The sulfur atoms of C105, C129, C131, and C151 are shown as yellow sticks, and the coordinated Zn²⁺ ion is shown as a gray sphere.

Whereas the NS1D and NS1A CTDs share the central twisted β-sheet, their surrounding α-helices adopt markedly different arrangements. In NS1A, the three C-terminal α-helices are located on the same side of the β-sheet (Fig. 4C). Likewise, in NS1B, the two α-helices following the central β-sheet adopt an arrangement similar to that of NS1A and distinct from that observed in NS1D (Fig. 4D).

Most significantly, NS1D-CTD lacks the first β-strand and α-helix present at the N-terminus of NS1A and NS1B CTDs. Instead, this region forms an extended loop structure containing four cysteine residues, C105, C129, C131, and C151, that coordinate a metal ion in tetrahedral geometry (Fig. 4E). The metal ion was identified as zinc by X-ray fluorescence spectroscopy and wavelength-dependent anomalous scattering. X-ray fluorescence emission spectra showed distinct peaks corresponding to the Zn Kα and Kβ emission lines, with no detectable peaks from other metal ions (Fig. S2A). The presence of zinc in the crystal was further confirmed by a Zn K-edge fluorescence scan (Fig. S2B). To verify the location of the zinc ion within the structure, diffraction datasets were collected at SSRL beamline 12-2 at 9,613 and 9,713 eV, immediately below and above the Zn K-edge, respectively. Anomalous difference maps calculated from these datasets revealed a pronounced change in peak height at the zinc-binding site, whereas peaks associated with sulfur atoms, used as internal controls, remained largely unchanged (Fig. S2C).

Although NS1C and NS1D share relatively low sequence identity (≤33.3%), phylogenetic and sequence alignment analyses revealed that all four cysteine residues involved in Zn²⁺ coordination are completely conserved across all nonredundant NS1C and NS1D sequences (Fig. S3). Notably, these four cysteine residues and the corresponding Zn²⁺-binding motif are absent from NS1A and NS1B, indicating that the Cys₄:Zn cluster is a distinctive structural feature of influenza C and D virus NS1 proteins.

To determine whether the Zn²⁺ ion contributes to NS1D-CTD folding and stability, we performed thermal shift assays in the presence or absence of 5 mM ethylenediaminetetraacetic acid (EDTA). Chelation of Zn²⁺ by EDTA significantly reduced the melting temperature of NS1D-CTD from 59.0 °C to 55.5 °C (Fig. S2D–E), indicating that the Cys4:Zn cluster contributes substantially to protein stability. Consistent with this observation, recombinant NS1D C105A and C129A mutants showed strong aggregation tendency and could not be purified.

### NS1D-CTD preferentially binds ssRNA through a conserved basic surface

During purification of NS1D-CTD, we observed a strong tendency of the protein to associate with *E. coli*-derived RNA. We therefore examined its RNA-binding activity by fluorescence polarization (FP) assays using the same 21-nt fluorescein-labeled RNA oligonucleotides used for NS1D-NTD. NS1D-CTD bound ssRNA with moderate affinity (K_D_ = 1,800 ± 50 nM) and dsRNA with slightly lower affinity (K_D_ = 3,135 ± 33 nM) (Fig. 5A). Interestingly, the crystallizable NS1D-CTD ΔCTT construct lacking the C-terminal tail exhibited enhanced ssRNA binding (*K*_D_ = 450 ± 13 nM) (Fig. 5B). To assess CTD-mediated RNA binding in the context of full-length NS1D, we tested an NS1D K37A/K40A mutant in which NTD-mediated RNA binding was disrupted. This mutant preferably bound ssRNA with approximately 7.5-fold higher affinity for ssRNA than dsRNA (K_D_ = 119 ± 7 nM vs. 900 ± 5 nM) (Fig. 5A). In contrast, no measurable FP signal was observed for H6N6 NS1A binding to ssRNA (Fig. 5C), consistent with previous reports [32].

**Figure 5.**
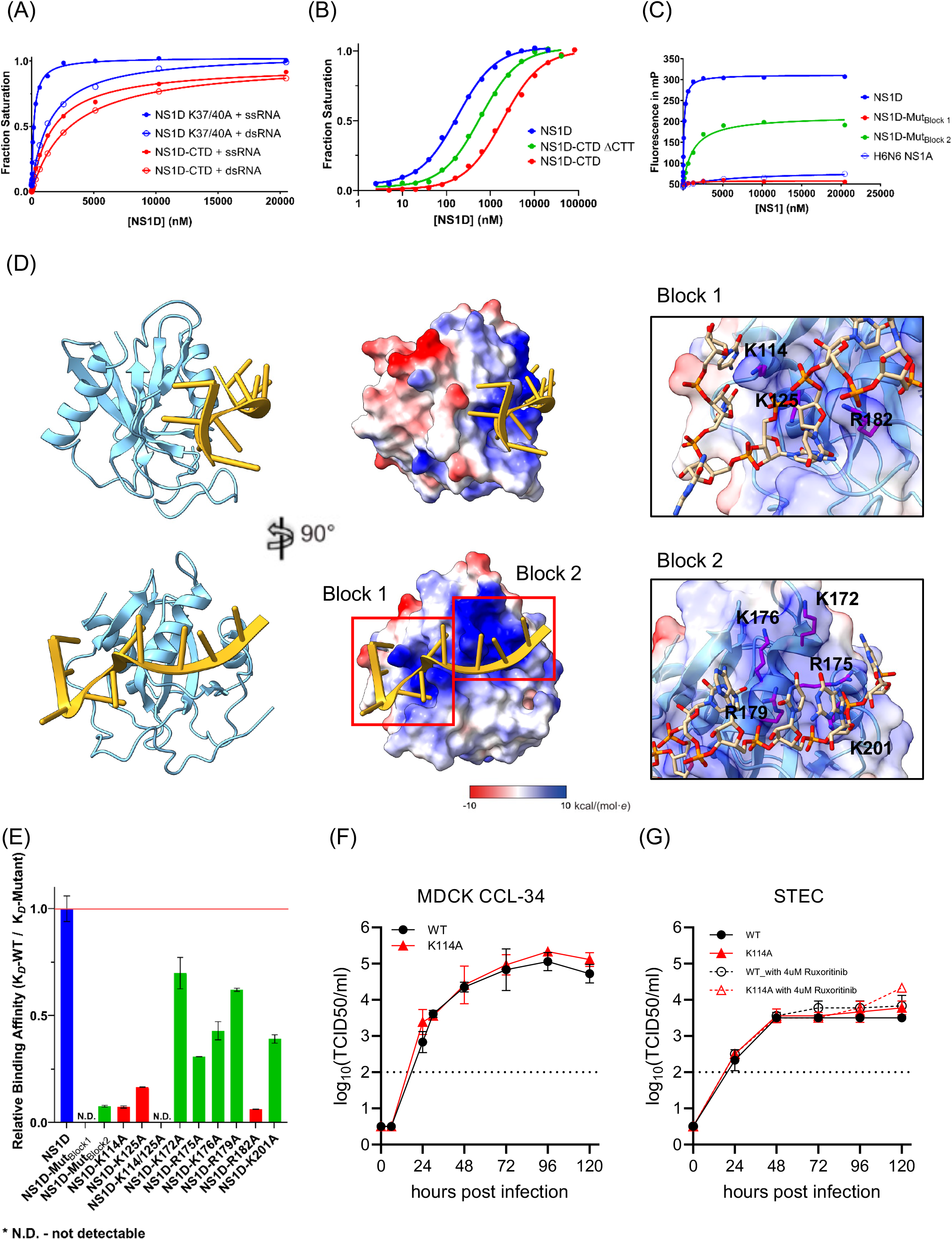
Molecular basis of NS1D-CTD–RNA binding. (A) Fluorescence polarization analysis of ssRNA and dsRNA binding by NS1D-CTD and full-length NS1D. (B) Comparison of ssRNA binding by NS1D and its truncations using fluorescence polarization assays. (C) Comparison of ssRNA binding by NS1D and H6N6 NS1A. (D) Molecular dynamics model of the NS1D-CTD–ssRNA complex. NS1D-CTD and ssRNA are shown in light sky blue and gold, respectively. The electrostatic surface potential of NS1D-CTD is shown with positively and negatively charged regions in blue and red, respectively. A positively charged surface centered on the α4 helix comprises two regions, block 1 and block 2, containing a total of eight Lys and Arg residues implicated in RNA binding. (E) Comparison of ssRNA binding by NS1D and the indicated mutants. (F, G) Growth kinetics of wild-type (WT) and K114A mutant viruses in MDCK CCL-34 and swine tracheal epithelial cells (STEC), in the presence or absence of 4 μM ruxolitinib. Cells were infected at a multiplicity of infection (MOI) of 0.005 PFU/cell. Error bars represent the standard deviation of triplicate measurements. The dotted line indicates the limit of detection (2 log₁₀ TCID₅₀/mL).

Because NS1D-NTD showed no detectable ssRNA binding (Fig. 3B), we next introduced mutations into full-length NS1D to identify residues within the CTD that contribute to ssRNA recognition. To guide the selection of residues for mutational analysis, we modeled the NS1D-CTD–RNA complex using HDOCK [39], assuming rigid-body docking without major conformational changes in either the protein or RNA upon complex formation. The resulting docking model was subjected to energy minimization using the standard CHARMM-GUI equilibration protocol [40] in NAMD 3.01 [41]. In the model, the 10-nt ssRNA was positioned along a positively charged surface centered on helix α4 (Fig. 5D). This surface could be spatially divided into two basic regions. Block 1 comprises a deep pocket formed by K114 in β1, K125 in β2, and R182 following α5, whereas block 2 comprises K172, R175, K176, and R179 in α4 together with K201 following α5.

We then performed FP assays on block mutants using the same fluorescein-labeled ssRNA oligonucleotide. Substitution of the five positively charged residues in block 2 reduced ssRNA-binding affinity by approximately 13-fold (*K*_D_ = 1,560 ± 86 nM), whereas substitution of the three positively charged residues in block 1 completely abolished RNA binding (Fig. 5E). Further analysis of individual residues within these basic regions showed that K114, K125, and R182 made the largest contributions to ssRNA binding. Consistent with these results, the K114A/K125A double mutant completely abolished detectable RNA-binding activity (Fig. 5E). Phylogenetic and sequence alignment analyses of NS1C and NS1D revealed strict conservation of the positively charged residues forming this RNA-binding surface, with the exception of K114, which is not conserved as a basic residue in NS1C (Fig. S3). This high degree of conservation suggests that NS1C and NS1D likely employ a similar basic surface for RNA recognition.

### The Zn²⁺-binding motif and conserved CTD basic surface are critical for recovery of infectious IDV

Based on our *in vitro* characterization of NS1D-CTD, we next investigated the contributions of Zn²⁺ coordination and the CTD RNA-binding surface to IDV replication using reverse genetics [42]. We introduced six mutations into the IDV NS segment to selectively disrupt either the Cys4:Zn cluster (C105A, C129A, and C131A) or the positively charged RNA-binding surface (K114A, K125A, and the K114A/K125A double mutant). These mutations are located within the intronic region of the NS2D transcript and are therefore not expected to directly alter the NS2D protein sequence.

After multiple virus rescue attempts, we recovered only WT IDV and the NS1D K114A mutant (Fig. 5F and G). Despite successful transfection, none of the three Cys4:Zn cluster mutants, the K125A mutant, or the K114A/K125A double mutant yielded recoverable infectious virus in cell culture, with HA titers remaining below 2 (Fig. S4). Similar results were obtained in the presence of 4 μM ruxolitinib, a JAK–STAT pathway inhibitor that suppresses interferon signaling and has been reported to enhance influenza virus replication in MDCK cells [43]. The inability to recover the Cys4:Zn cluster mutants is consistent with the structural role for Zn²⁺ coordination in maintaining CTD stability, as suggested by our thermal stability measurements and the aggregation observed for recombinant C105A and C129A proteins (Fig. S2D–E). K125, which is strictly conserved among NS1D and NS1C sequences, lies within the positively charged RNA-binding surface of NS1D-CTD and was identified by our *in vitro* assays as a key determinant of RNA binding. The failure to recover the K125A and K114A/K125A mutant viruses therefore suggests that K125 contributes to an important function required for productive IDV infection, potentially involving RNA recognition or other molecular interactions mediated by the CTD.

In contrast, the NS1D K114A mutant was successfully rescued, although its initial HA titers were slightly lower than those of WT virus (Fig. S4B, C). Following recovery, the K114A virus exhibited replication kinetics comparable to those of WT virus in both MDCK cells and swine tracheal epithelial cells (STECs) (Fig. 5F–G). Multistep growth experiments performed at an MOI of 0.005 revealed no significant differences between WT and K114A viruses under the conditions tested, including with treatment or without treatment of 4 μM ruxolitinib.

Together, these findings indicate that residues involved in Zn²⁺ coordination and the conserved CTD basic surface are important for recovery of infectious IDV. The contrasting phenotypes of K114A and K125A further suggest that the biological contributions of individual CTD residues cannot be explained solely by their effects on RNA-binding affinity measured in vitro.

## Discussion

While NS1D retains the general two-domain organization of influenza virus NS1 proteins, both domains exhibit features that distinguish NS1D from the more extensively characterized NS1 proteins of influenza A and B viruses. In particular, NS1D-CTD contains a previously unrecognized Zn²⁺-binding motif and a distinct RNA-binding surface, highlighting structural and functional diversity among influenza virus NS1 proteins.

Although the NS1D-NTD (residues 1-77) adopts the dimeric six-helical fold characteristic of influenza virus NS1 proteins, but the C-terminal α3 helix of each NS1D protomer is considerably shorter than its counterparts in NS1A and NS1B, resulting in a longer, potentially more flexible linker between the NTD and CTD (Fig. 2). This increased flexibility may influence the relative disposition of the CTDs within the full-length NS1D dimer and thereby contribute to functional differences. Structural modeling of the NS1D-NTD-dsRNA complex identified several basic residues, including R13, K37, R39, and K40, positioned near the phosphate backbone of dsRNA. Functional analysis supported the importance of this conserved basic region, as mutation of K37 and K40 abolished detectable dsRNA binding. Notably, NS1D-NTD showed a clear preference for dsRNA, with no detectable binding to ssRNA under the same assay conditions, suggesting that the NTD is specialized for dsRNA recognition rather than functioning as a general RNA-binding domain. Together, these observations indicate that NS1D preserves a conserved mode of dsRNA recognition while exhibiting structural adaptations that may distinguish its function from those of NS1A and NS1B [15].

In contrast, NS1D-CTD (residues 90–231) adopts a substantially different structural arrangement from the CTDs of NS1A and NS1B. Although the central five-stranded β-sheet is conserved, the surrounding α-helices differ markedly in their positioning, resulting in a distinct overall topology. In addition, the N-terminal region of NS1D-CTD forms an elaborate loop containing four cysteine residues that tetrahedrally coordinate a Zn²⁺ ion. This Cys4:Zn cluster is absent from known NS1A and NS1B structures and does not appear to resemble commonly observed structural zinc-binding motifs [16,38].

Functionally, NS1D-CTD preferentially binds ssRNA over dsRNA, and mutational analysis identified a conserved positively charged surface that likely mediates RNA recognition. This RNA-binding profile differs from those of NS1A- and NS1B-CTDs. NS1A showed no detectable binding to ssRNA under our assay conditions, consistent with previous reports [32] (Fig. 5). NS1B-CTD has been reported to preferentially bind dsRNA over ssRNA [16], with particularly strong binding to 5′-triphosphorylated dsRNA [44]. Another notable difference concerns oligomerization. Whereas NS1A- and NS1B-CTDs can form dimeric assemblies [16,38], we observed no evidence of NS1D-CTD dimerization either in the crystal structure or in solution. Thus, our findings indicate that NS1D-CTD has evolved distinct structural, metal-binding, oligomerization, and RNA-binding properties.

Phylogenetic analyses have shown that the NS segment displays the greatest evolutionary divergence among the genome segments of the four influenza virus genera [17]. Consistent with this observation, NS1D shares only approximately 30% identity with NS1C and even lower identity with NS1A and NS1B. Our phylogenetic analysis of all nonredundant NS1C and NS1D sequences likewise revealed that the two proteins form distinct clades, reflecting substantial sequence divergence between ICV and IDV (Fig. S3). Remarkably, however, this extensive divergence contrasts with strong conservation of key structural and functional features within the CTD. The four cysteine residues that coordinate the structural Zn²⁺ ion are invariant across NS1C and NS1D sequences, while the basic character of the RNA-binding surface is also highly conserved, with seven of the eight strictly conserved basic residues in the CTD mapped to this area (i.e. K125, K172, R175, K176, R179, R182 and K201). Although direct structural and biochemical characterization of NS1C will be required to establish whether these residues mediate similar functions, their conservation suggests that NS1C may possess a comparable zinc-binding architecture and RNA-interaction surface. Together with our biochemical, mutational, and virus-rescue results, these observations support important roles for Zn²⁺-dependent stabilization of the CTD and CTD-mediated RNA recognition in NS1D function and further suggest that these structural and functional features may also be conserved in NS1C.

A particularly striking feature of NS1D is the apparent requirement for an intact CTD for IDV replication. This contrasts with IAV, for which viruses lacking NS1 or containing substantial NS1 truncations can be rescued when NEP/NS2 expression is preserved, although such viruses are strongly attenuated and often require permissive, interferon-deficient conditions for efficient propagation [1,43]. In our IDV system, by contrast, even single-residue substitutions disrupting either the structural Cys4:Zn cluster (C105A, C129A, or C131A) or the conserved CTD basic surface (K125A) were sufficient to prevent recovery of infectious virus. The inability to rescue these point mutants even in the presence of ruxolitinib, a JAK–STAT pathway inhibitor, suggests that the defects associated with these mutations cannot be readily overcome by chemical inhibitor suppression of JAK–STAT signaling. However, the inability to recover these mutants does not establish the precise molecular basis of their defects. Disruption of Zn²⁺ coordination may compromise CTD folding or stability, whereas substitution of K125 may affect RNA binding or other molecular interactions. In addition, although these mutations do not directly alter the NS2D protein sequence, potential effects on NS-segment RNA processing or NS2D expression cannot be excluded. Further studies will therefore be required to determine the molecular mechanisms underlying these phenotypes.

The contrasting phenotypes of K114 and K125 also indicate that the biological function of CTD is likely more complex than can be inferred from RNA-binding measurements alone. Although both residues contributed substantially to ssRNA binding *in vitro*, K125A could not be rescued, whereas K114A remained viable and displayed replication kinetics comparable to WT virus in both MDCK cells and STECs. Thus, the contribution of CTD to IDV replication does not correlate simply with its RNA-binding activity measured using recombinant protein. Consistent with this distinction, K114 is not conserved in NS1C, where it is replaced by a threonine, suggesting that a basic side chain at this position is not required for a conserved function of the CTD. In contrast, the strong conservation and virus-rescue phenotype of K125 suggest a more fundamental role that may extend beyond RNA binding, potentially involving additional viral or host interactions. Further characterization of this surface will be required to define its molecular partners and clarify its contributions to viral replication and antagonism of host antiviral responses.

Although we determined high-resolution crystal structures of the NS1D-NTD and NS1D-CTD, the conformations of the 12-residue linker region (LR) and 12-residue CTT remain unresolved. Crystals of NS1D-NTD-LR (residues 1–88) showed no interpretable electron density for the LR, whereas NS1D-LR-CTD (residues 78–231) and NS1D-CTD-CTT (residues 90–243) did not yield diffraction-quality crystals. To gain insight into the organization of full-length NS1D, we used AlphaFold3 [45] to model the orientation of the CTDs and the connectivity provided by the LR. The top-ranked AlphaFold3 model adopted an open dimeric conformation resembling that previously described for H5N1 NS1 (PDB ID: 3F5T) [13] (Fig. 6A). When our energy-minimized NS1D-NTD–dsRNA and NS1D-CTD–ssRNA models were superimposed onto the corresponding domains of the full-length AlphaFold3 [45] model, both RNA molecules could be accommodated simultaneously without steric clashes (Fig. 6B). Although this model remains to be experimentally validated, it raises the possibility that full-length NS1D could engage distinct RNA species through its NTD and CTD simultaneously or independently, with the flexible LR allowing substantial variation in the relative orientation of the two domains.

**Figure 6.**
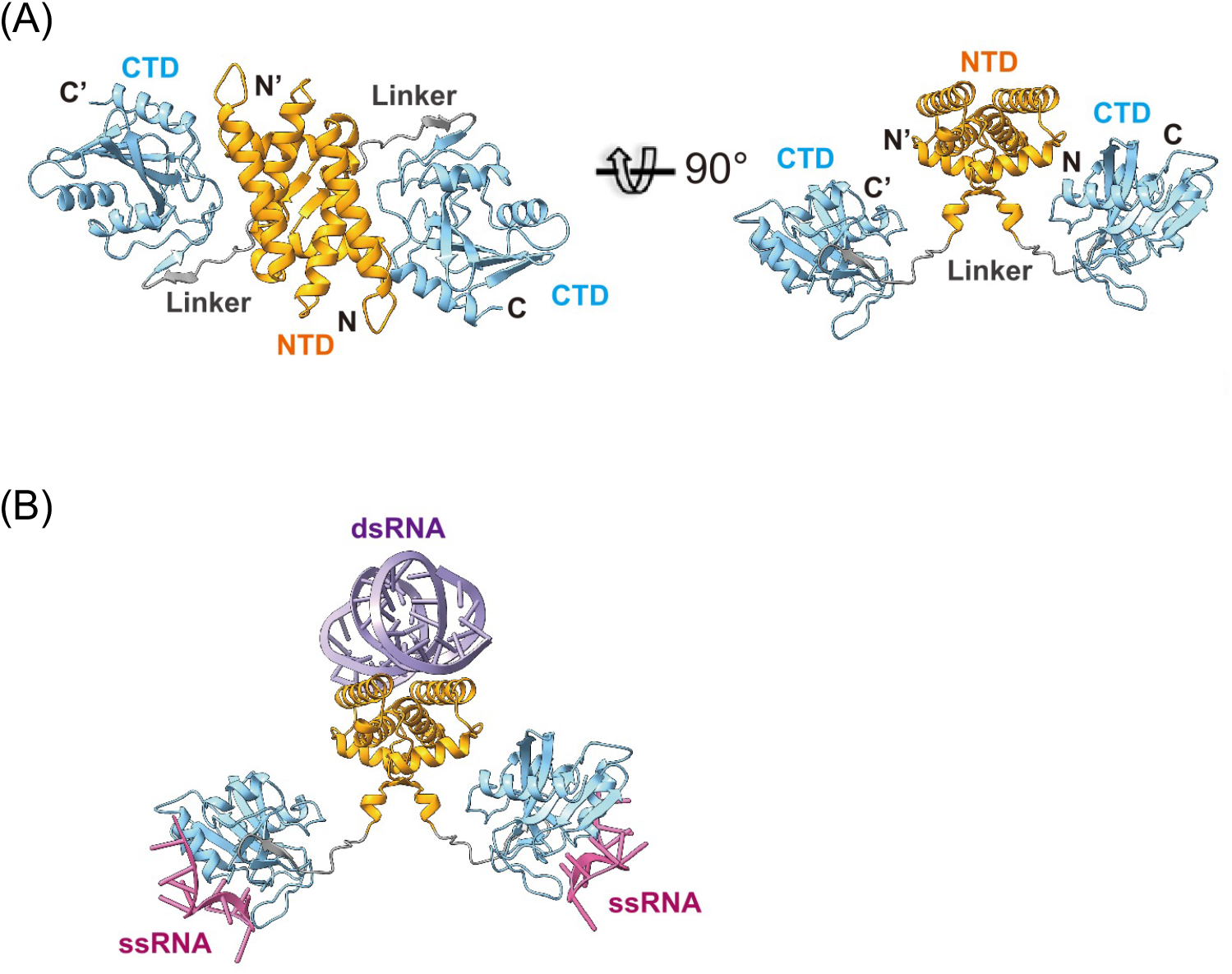
Structural modeling of RNA binding by full-length NS1D. (A) Structural model of full-length NS1D. The model illustrates the spatial arrangement of the NTD and CTD, which are connected by the linker region (LR; residues 78–89). Ribbon representations of the NTD, LR, and CTD are colored orange, gray, and light sky blue, respectively. (B) Structural model of full-length NS1D simultaneously bound to dsRNA and ssRNA. The dsRNA and ssRNA molecules are shown in purple and red, respectively. The model highlights the LR flexibility and suggests that full-length NS1D can accommodate dsRNA binding through the NTD and ssRNA binding through the CTD simultaneously without steric clashes.

This model provides a possible explanation for the enhanced ssRNA-binding affinity of full-length NS1D compared with NS1D-CTD alone (Fig. 5A), as flexibility within the LR may allow both CTDs of the NS1D dimer to engage the ssRNA substrate simultaneously. In addition, NS1D-CTD ΔCTT, which lacks the C-terminal 12 residues, bound ssRNA more strongly than NS1D-CTD (Fig. 5B), suggesting that the CTT may modulate RNA binding. The CTT sequence (IQAGSSDGELFD), which was not present in our crystallization construct, contains several negatively charged residues that could modulate electrostatic interactions between the CTD and RNA.

In summary, our results show that NS1D retains the conserved domain architecture of influenza virus NS1 proteins while exhibiting distinct structural and functional properties, particularly within the CTD. The Cys₄:Zn cluster and conserved basic RNA-binding surface of the CTD are associated with functions important for recovery of infectious IDV, and the conservation of these features in NS1C suggests that they may represent shared functional elements of influenza C and D virus NS1 proteins. These findings provide a structural framework for understanding NS1D function, suggest that NS1D-CTD may contribute to multiple aspects of viral infection beyond its established role in immune antagonism, and further distinguish NS1D from its more extensively characterized influenza A and B virus counterparts.

## Materials and Methods

### Plasmid construction

Influenza D virus (IDV) strain D/bovine/Mississippi/C00180N/2015 (C00180N) was propagated in Madin-Darby canine kidney (MDCK) cells. The NS1D coding sequence was amplified as cDNA generated by reverse transcription of RNA isolated from infected cells. The coding sequence corresponding to the NS1D N-terminal domain (NTD) was cloned into the pETDuet-His-SUMO expression vector, whereas the full-length NS1D coding sequence was cloned into the pET28A-His-Trx-TEV vector. The pET28A-His-Trx-TEV-NS1D construct was used to generate the K37A/K40A and Δ232–243 mutations.

### Protein expression and purification

The pETDuet-His-SUMO-NS1D-NTD construct was expressed in *Escherichia coli* BL21 (DE3) cells. Cultures were grown at 37 °C in M9 minimal medium supplemented with selenomethionine (Se-Met; TCI America) to an OD_600_ of 0.8, induced with 0.2 mM isopropyl β-D-1-thiogalactopyranoside (IPTG), and incubated overnight at 16 °C. Cell pellets were resuspended in lysis buffer A (50 mM Tris-HCl, pH 8.0, 150 mM NaCl, 1 mM NaN_3_, and 5 mM β-mercaptoethanol) and lysed by sonication. The clarified lysate was applied to a 5-mL HisTrap column (Cytiva), washed with lysis buffer A, and eluted with a linear imidazole gradient to a final concentration of 1 M. The His-SUMO tag was removed by incubation with yeast Ulp1 protease, and the cleaved NS1D-NTD was further purified by HiTrap Heparin HP and HiLoad 16/600 Superdex 75 pg chromatography (Cytiva). The purified protein was concentrated to 10 mg/mL in lysis buffer A.

The pET28A-His-Trx-TEV-NS1D K37A/K40A Δ232–243 construct was similarly expressed in *E. coli* BL21(DE3) cells. Cell pellets were resuspended in lysis buffer B (50 mM Tris-HCl, pH 8.0, 500 mM NaCl, 500 mM urea, 1 mM NaN_3_, and 1 mM dithiothreitol [DTT]) and lysed by sonication. The clarified lysate was subjected to nickel-affinity chromatography using HisPur Ni-NTA resin (Thermo Fisher Scientific). After three washes with wash buffer (50 mM Tris-HCl, pH 8.0, 500 mM NaCl, 30 mM imidazole, 1 mM NaN_3_, and 1 mM DTT), the resin was resuspended in 20 mL of wash buffer and incubated overnight at 4 °C with 0.05 mg/mL trypsin to release NS1D-CTD (residues 90–231). Digestion was terminated by the addition of phenylmethylsulfonyl fluoride (PMSF), after which the reaction mixture was diluted twofold with low-salt buffer (50 mM Tris-HCl, pH 8.0, 1 mM NaN_3_, and 1 mM DTT) and further purified by HiTrap Heparin HP chromatography to remove residual trypsin. Purified NS1D-CTD was subsequently subjected to size-exclusion chromatography on a Superdex 200 Increase 10/300 GL column (Cytiva) and concentrated to 5 mg/mL in buffer containing 50 mM Tris-HCl (pH 8.0), 300 mM NaCl, 200 mM KCl, 1 mM NaN_3_, and 1 mM DTT.

### Crystallization and data collection

Crystals of Se-Met-derivatized NS1D-NTD residues 1–77 were obtained by hanging-drop vapor diffusion by mixing protein and reservoir solution at a 1:1 ratio. The reservoir solution contained 100 mM sodium acetate (pH 7.0), 1.2 M ammonium sulfate, and 200 mM lithium sulfate. Crystals were transferred to cryoprotectant solution containing reservoir solution supplemented with 25% glycerol and flash-frozen in liquid nitrogen. Diffraction data were collected at Advanced Photon Source beamline 21-ID-G using a MAR 300-mm detector.

Crystals of NS1D-CTD were obtained by hanging-drop vapor diffusion by mixing protein and reservoir solution at a 1:1 ratio. The reservoir solution contained 100 mM HEPES (pH 7.5) and 10% (w/v) PEG 3350. Crystals were transferred to cryoprotectant solution containing reservoir solution supplemented with 25% glycerol and flash-frozen in liquid nitrogen. Diffraction data used to generate the initial electron-density model of NS1D-CTD were collected using a Rigaku XtaLAB Synergy-S-DW diffractometer equipped with a HyPix-Arc 150° hybrid photon-counting detector and a Cu microfocus sealed X-ray source operating at a wavelength of 1.5406 Å. Higher-resolution diffraction data were collected at the Stanford Synchrotron Radiation Lightsource. Data were acquired at beamline 12-2 using a Dectris EIGER 16M PAD detector at multiple wavelengths and at beamline 14-1 using a Dectris Pilatus 6M detector at a wavelength of 1.19591 Å.

### Data processing and structural determination

Diffraction data from Se-Met-derivatized NS1D-NTD crystals were indexed, integrated, and scaled using HKL2000 [46]. The crystals belonged to space group C2, and data collection statistics are summarized in Table 1. Initial phases were determined using the Phenix AutoSol Wizard [47], followed by iterative cycles of model building and refinement in COOT [48] and Phenix.refine [49].

Diffraction data from the initial NS1D-CTD crystals were processed using XDS [50]. The crystals belonged to space group P2₁2₁2₁, and data collection statistics are summarized in Table 1. An AlphaFold2 [36] model of NS1D-CTD residues 90–231 was generated using ColabFold [37] with 12 recycling iterations to improve prediction consistency given the limited depth of the multiple sequence alignment. The relaxed top-ranked model was processed in PyMOL [51], and residues with pLDDT values greater than 80 that were assigned as α-helices or β-strands by the DSS algorithm were retained to generate a core search model. pLDDT scores were converted to B-factors using values of 1500/pLDDT for main-chain atoms and 2200/pLDDT for side-chain atoms, resulting in B-factors ranging from 16–20 Å² and 23–29 Å², respectively. Molecular replacement was performed with PHASER, yielding a solution with a TFZ score of 8.8 and an LLG score of 64. The initial model was further improved by density modification and automated model building using SHELXE [52] and ARP/wARP [53]. Following collection of higher-resolution diffraction data at SSRL, the final model was completed through iterative cycles of model building and refinement using COOT [48] and Phenix.refine [49].

All structural biology software used in this study was provided through the SBGrid consortium [54]. Structural figures were generated using ChimeraX [55].

### Fluorescence polarization assays

RNA oligonucleotides used for fluorescence polarization (FP) assays were purchased from Integrated DNA Technologies (Coralville, IA). Double-stranded RNA (dsRNA) was generated by annealing two complementary 21-nt single-stranded RNAs (ssRNAs), ACGACCUAAUUAUGCUUGAUG-AlexF488N (sense) and CAUCAAGCAUUAUUAGGUCGU (antisense), in hybridization buffer containing 50 mM Tris-HCl (pH 8.0), 150 mM NaCl, 1 mM NaN3, and 5 mM β-mercaptoethanol. The RNA mixture was heated to 95 °C for 2 min and gradually cooled to 25 °C over 45 min. The resulting dsRNA stock solution (3 µM) was diluted 100-fold in FP reaction buffer (20 mM HEPES, pH 7.4, 50 mM KCl, 0.1 mM MgCl_2_, and 0.5 mM DTT) to obtain a final concentration of 30 nM.

For ssRNA-binding assays, 5 µL of 30 nM fluorescein-labeled sense ssRNA was mixed with 20 µL of serially diluted NS1 proteins to determine the dissociation constants (K_D_). For dsRNA-binding assays, 5 µL of 30 nM dsRNA was incubated with 20 µL of serially diluted NS1 proteins. Protein samples were serially diluted twofold in FP reaction buffer to generate a broad concentration range. Fluorescence measurements were acquired after a 5-min incubation at room temperature using a Spark Multimode Microplate Reader (Tecan) in low-volume 384-well black/clear flat-bottom microplates (Corning). Binding curves were fitted by nonlinear regression using Microsoft Excel Solver and GraphPad Prism. All experiments were performed in triplicate, and the data are presented as the mean values.

### Thermal shift assays

Protein thermal shift assays (TSAs) were performed in triplicate using Protein Thermal Shift kits (Applied Biosystems) according to the manufacturer’s instructions. Reactions were conducted on a CFX96 Touch Real-Time PCR Detection System (Bio-Rad) and analyzed using CFX Manager software (Bio-Rad). Each 25-µL reaction contained 20 mM Tris-HCl (pH 8.0), 100 mM NaCl, and 25 µg of NS1D-CTD (residues 90–231), in the presence or absence of 5 mM ethylenediaminetetraacetic acid (EDTA). No-protein controls were included for all conditions. Samples were heated from 25 °C to 99 °C at a rate of 0.05 °C/s, and fluorescence measurements were recorded every 0.5 °C. Relative fluorescence units (RFU) and derivative melting curves (dRFU/dT) were plotted using GraphPad Prism.

### Molecular dynamics simulations

The ssRNA sequence used for molecular docking (CGUAUUCGUC) was generated by HDOCK homology modeling [39] using a single-strand RNA molecule from PDB ID: 6N8I as the template [56]. HDOCK generated 100 docking models that were ranked according to docking and confidence scores. The top 10 models are presented in Figure S5 and Table S1. Docking model 10 was selected because it was consistent with independent AlphaFold3 [45] predictions using five different RNA oligo sequences that exhibited high ipTM and pTM scores (Fig. S6). The docking model was further refined by energy minimization and molecular dynamics simulations.

All-atom molecular dynamics (MD) simulations were performed to investigate interactions between NS1D-CTD and ssRNA. Simulation systems were prepared using the CHARMM-GUI Solution Builder [40] with a 2-nm solvent buffer in all directions, yielding a simulation box of 8.4 × 8.4 × 8.4 nm³. The system was neutralized and adjusted to 50 mM KCl at pH 7.4.

Following standard CHARMM-GUI equilibration procedures, production simulations were performed in NAMD 3.01 [41] using the CHARMM36m force field [57] and the TIP3P water model with a 2-fs integration time step. Simulations were conducted in the NVT ensemble at 294 K. Lennard-Jones interactions were truncated at 12 Å with a switching function beginning at 10 Å, and long-range electrostatic interactions were calculated using the particle mesh Ewald (PME) method [58] with a 12-Å cutoff. Periodic boundary conditions were applied throughout the simulations. The following NAMD3 settings were used: CUDASOAintegrate = on, margin = 8, stepsPerCycle = 400, and pairlistsPerCycle = 40. Independent trajectories were generated for subsequent analyses.

All simulations were performed on the Carya cluster at the Data Science Institute, University of Houston.

### Phylogenetic analysis

Influenza C and D virus NS1 protein sequences were downloaded from the NCBI Protein database [59] on September 4, 2024. Sequences were aligned using the T-Coffee Expresso structural alignment tool [60]. The resulting alignment was curated in Jalview [61] to remove redundant and partial sequences, followed by additional manual inspection. Branch support was calculated using ultrafast bootstrap analysis on the W-IQ-TREE web server [62], with 1,000 bootstrap replicates and 1,000 iterations. The phylogenetic tree was visualized using iTOL v6 [63], colored according to the Clustal scheme, and displayed with a 100% consensus threshold.

### Cells, virus rescue, and virus propagation

Madin-Darby canine kidney (MDCK; ATCC CCL-34), HEK293T (ATCC CRL-3216), and human rectal tumor (HRT-18G; ATCC CRL-3609) cells were obtained from ATCC. Swine tracheal epithelial (STE) cells were kindly provided by Stacey Schultz-Cherry (St. Jude Children’s Research Hospital). All cell lines were maintained in Dulbecco’s modified Eagle’s medium (DMEM; Thermo Fisher Scientific) supplemented with 10% fetal bovine serum (FBS) at 37 °C in a humidified incubator with 5% CO₂.

The seven genomic segments of D/bovine/Oklahoma/660/2013 (D/OK) cloned into the pHW2000 vector were kindly provided by Dr. Feng Li (University of Kentucky). NS mutants (K114A, K125A, and K114A/K125A) were generated by site-directed mutagenesis (Gene Universal, Newark, DE). Recombinant viruses were rescued using a plasmid-based reverse genetics system as previously described [42]. Briefly, plasmids encoding the six non-NS genomic segments together with either wild-type or mutant NS segments were co-transfected into cocultured MDCK and HEK293T cells in 6-well plates using TransIT-LT1 reagent (Mirus) according to the manufacturer’s instructions. At 24 h post-transfection, TPCK-treated trypsin was added to a final concentration of 1 µg/mL. Supernatants were harvested at 96 h post-transfection and subsequently propagated in MDCK and HRT-18G cells. The NS segments of all rescued viruses were verified by Sanger sequencing to confirm the absence of unintended mutations.

### Multistep growth kinetics

Multistep growth kinetics were performed in MDCK and STE cells to evaluate the effects of NS mutations on viral replication. Cells were seeded in 6-well plates and cultured for approximately 18 h to reach ∼90% confluence. Monolayers were infected at a multiplicity of infection (MOI) of 0.005. Following virus adsorption, inocula were removed, cells were washed, and infection medium supplemented with 1 µg/mL TPCK-treated trypsin was added. For STE cells, parallel infections were performed in the presence of 4 µM ruxolitinib [43], a JAK1/2 inhibitor, to suppress JAK–STAT signaling. Supernatants were collected at 0, 24, 48, 72, 96, and 120 h post-infection (hpi), stored at −80 °C, and subsequently subjected to viral titration. All experiments were performed with three independent biological replicates.

### Viral titration and hemagglutination assays

Virus titers were determined by the 50% tissue culture infectious dose (TCID₅₀) assay in MDCK cells. Cells were seeded at 2 × 10 cells per well in 96-well plates containing Opti-MEM I reduced-serum medium (Thermo Fisher Scientific) and incubated for 18–20 h at 37 °C in 5% CO₂. Virus samples were serially diluted 10-fold in Opti-MEM supplemented with 1 µg/mL TPCK-treated trypsin, and 100 µL of each dilution was added to quadruplicate wells. Plates were incubated for 120 h, after which infection was determined by hemagglutination assay. TCID₅₀ values were calculated using the Reed–Muench method [64].

Hemagglutination assays were performed using 0.5% turkey erythrocytes (LAMPIRE Biological Laboratories) as previously described [65]. Briefly, serially diluted virus samples were incubated with erythrocytes, and hemagglutination activity was recorded to determine the presence or absence of infection in each well for TCID₅₀ calculations.

## Acknowledgements

This work was supported by Welch Foundation (C-1565 to YJT). This research used resources of the Advanced Photon Source, a U.S. Department of Energy (DOE) Office of Science User Facility operated for the DOE Office of Science by Argonne National Laboratory under Contract No. DE-AC02-06CH11357. Use of the LS-CAT Sector 21 was supported by the Michigan Economic Development Corporation and the Michigan Technology Tri-Corridor (Grant 085P1000817). Use of the Stanford Synchrotron Radiation Lightsource, SLAC National Accelerator Laboratory, is supported by the U.S. Department of Energy, Office of Science, Office of Basic Energy Sciences under Contract No. DE-AC02-76SF00515. The SSRL Structural Molecular Biology Program is supported by the DOE Office of Biological and Environmental Research, and by the National Institutes of Health, National Institute of General Medical Sciences (P30GM133894).

## Data availability

The structural coordinates have been deposited in the Protein Data Bank (PDB) under the accession number of 39II and 39IH for NS1D-NTD (Se-Met) and NS1D-CTD crystal structures, respectively.

