## Supplementary Information (PDF) for "Structure of influenza D virus NS1 reveals a novel Zn-binding site and ssRNA-binding activity"

### Supporting Information

**S1 Fig. Secondary structure assignments of NS1D generated using ChimeraX.**

**S2 Fig. Experimental identification of $Zn^{2+}$ as the metal ion coordinated by NS1D-CTD.**

**S3 Fig. Phylogenetic analysis and multiple sequence alignment of nonredundant NS1C and NS1D sequences.**

**S4 Fig. Hemagglutination (HA) titers from wild-type and NS1-mutant IDV rescue experiments.**

**S5 Fig. Ten highest-scoring docking models of the NS1D-CTD–ssRNA complex.**

**S6 Fig. Structural modeling of NS1D-CTD–ssRNA complexes.**

**S1 Table. Summary of the ten highest-scoring NS1D-CTD–ssRNA docking models.**

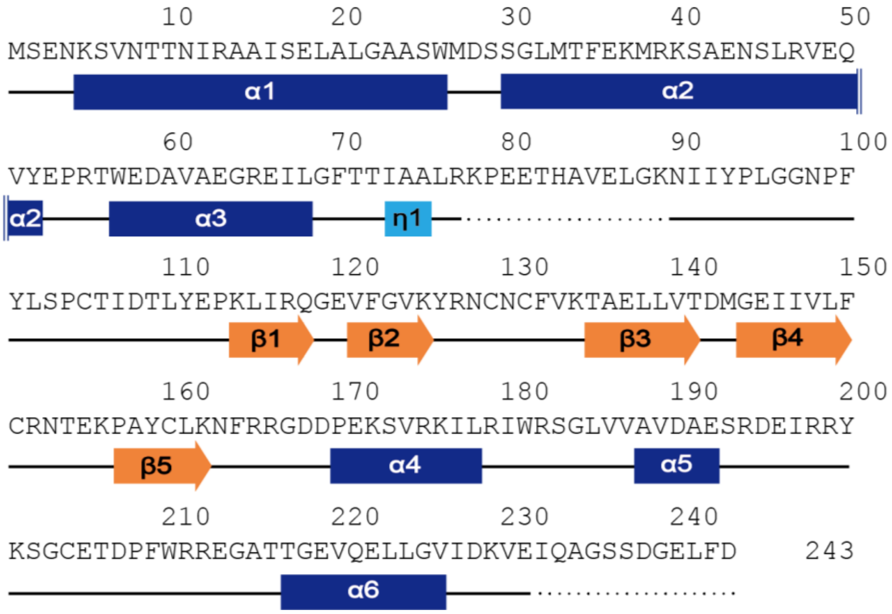


**S1 Fig. Secondary structure assignments of NS1D generated using ChimeraX.** α-Helices, β-strands, and loops are shown as rectangles, arrows, and thin lines, respectively. Unresolved regions are indicated by dotted lines.

(A) (B)

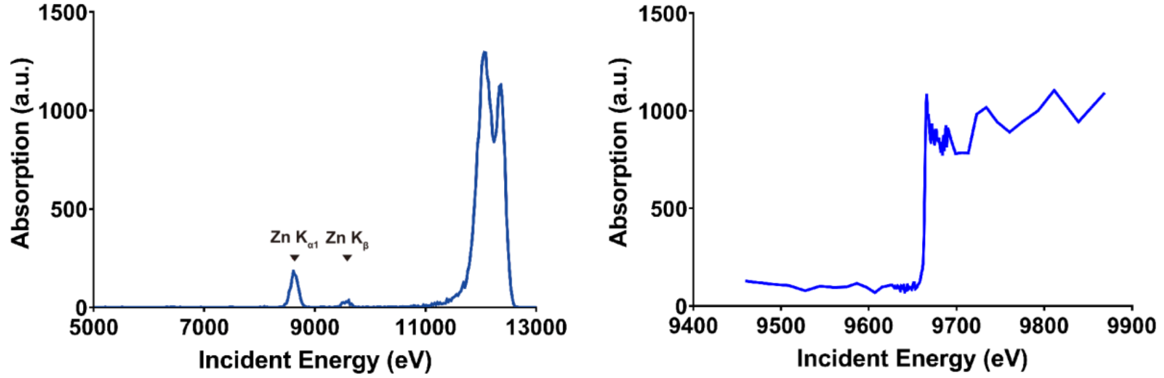


(C)

| Anomalous different Fourier peaks from the Zn atom | | | | | |
|---|---|---|---|---|---|
| | Zn-301 | Sulfur (Sγ) of Cys105 | Sulfur (Sγ) of Cys129 | Sulfur (Sγ) of Cys131 | Sulfur (Sγ) of Cys151 |
| λ2 above Zn (9713 eV) | 57.53 | 5.16 | 5.79 | 4.73 | 4.77 |
| λ3 below Zn (9613 eV) | 12 | 10.41 | 9.87 | 9.79 | 8.91 |

(D) (E)

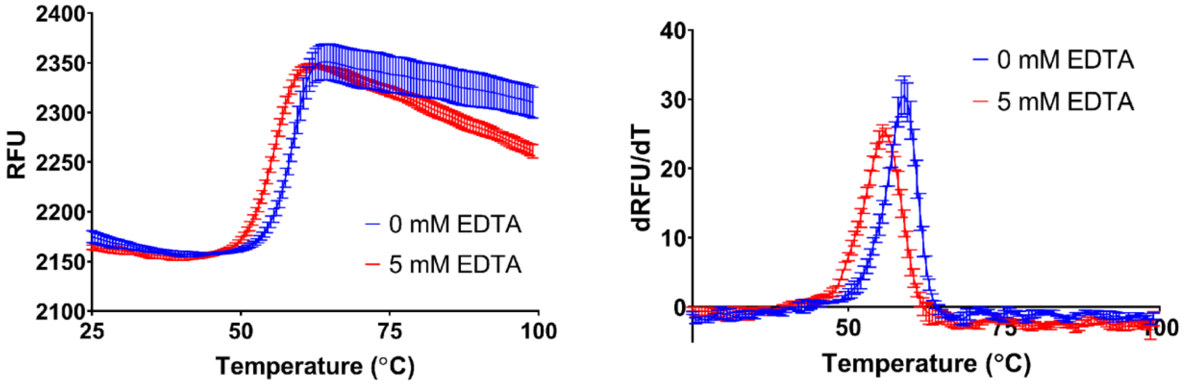


**S2 Fig. Experimental identification of $Zn^{2+}$ as the metal ion coordinated by NS1D-CTD.** (A–B) X-ray absorption spectroscopy (A) and Zn K-edge fluorescence scanning (B) confirmed zinc as the anomalous scatterer. (C) Anomalous difference electron density maps for Zn and the Sγ atoms of Cys105, Cys129, Cys131, and Cys151 collected at X-ray energies above and below the Zn absorption edge. (D) Thermal denaturation profiles of NS1D-CTD with and without EDTA treatment. (E) First-derivative melting curves (dRFU/dT) used to determine the melting temperature (Tm), represented by the peak of each curve. Chelation of Zn by EDTA reduced the thermal stability of NS1D-CTD, supporting a structural role for the Zn ion in maintaining domain integrity.

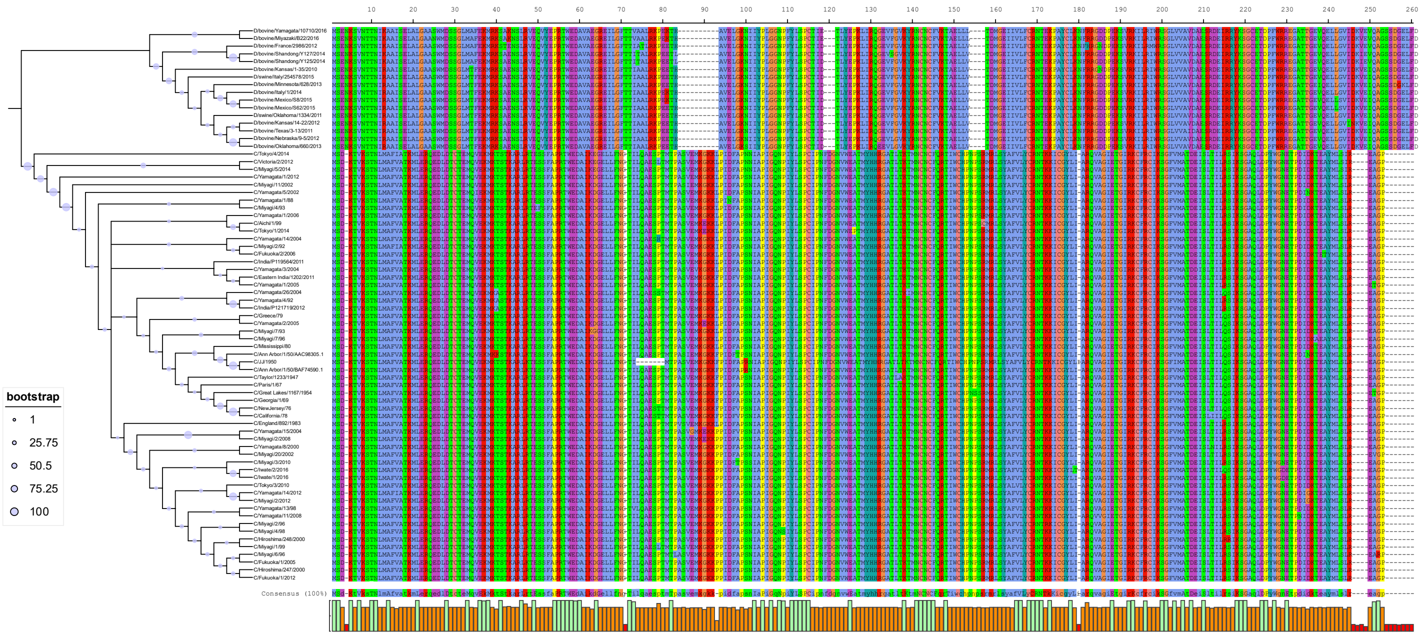

**S3 Fig. Phylogenetic analysis and multiple sequence alignment of nonredundant NS1C and NS1D sequences.** Key residues involved in formation of the $Cys_4$:Zn structural cluster and the RNA-binding pocket of NS1-CTD are invariant, displaying 100% conservation among all sequences analyzed.

(A)

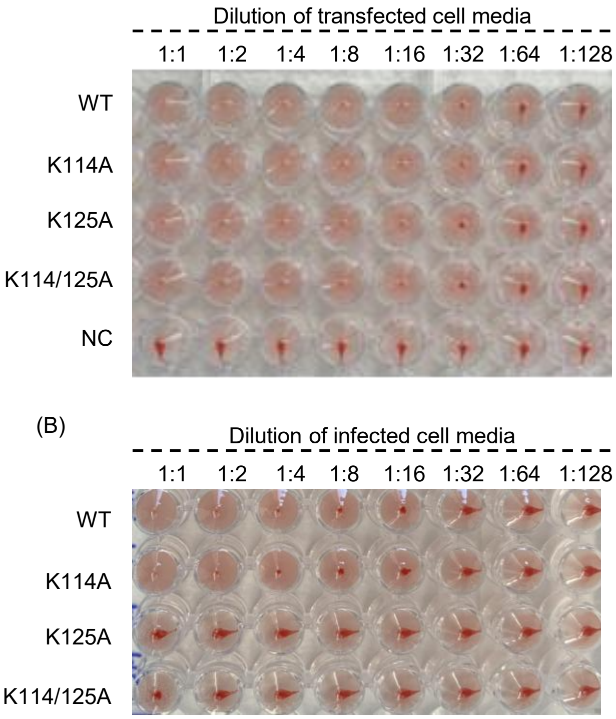


(C)

| Genotype | HA titer (turkey RBCs) |
|---|---|
| WT | 16 |
| C105A | <2 |
| K114A | 8 |
| K114/125A | <2 |
| K125A | <2 |
| C129A | <2 |
| C131A | <2 |

**S4 Fig. Hemagglutination (HA) titers from wild-type and NS1-mutant IDV rescue experiments.** (A) Representative HA titers of supernatants collected from transfected cells during virus rescue. Samples include wild-type (WT), NS1 mutants (K114A, K125A, and K114A/K125A), and a negative control (NC). (B) Representative HA titers obtained following infection of naïve cells with rescued virus preparations. Virus rescue was considered unsuccessful when HA titers remained below 2 after three serial passages of transfection supernatants in both MDCK and HRT-18G cells. (C) Summary of HA titers from successfully rescued NS1 mutant viruses following propagation in MDCK cells. Among the mutants tested, only the K114A mutant virus was successfully rescued.

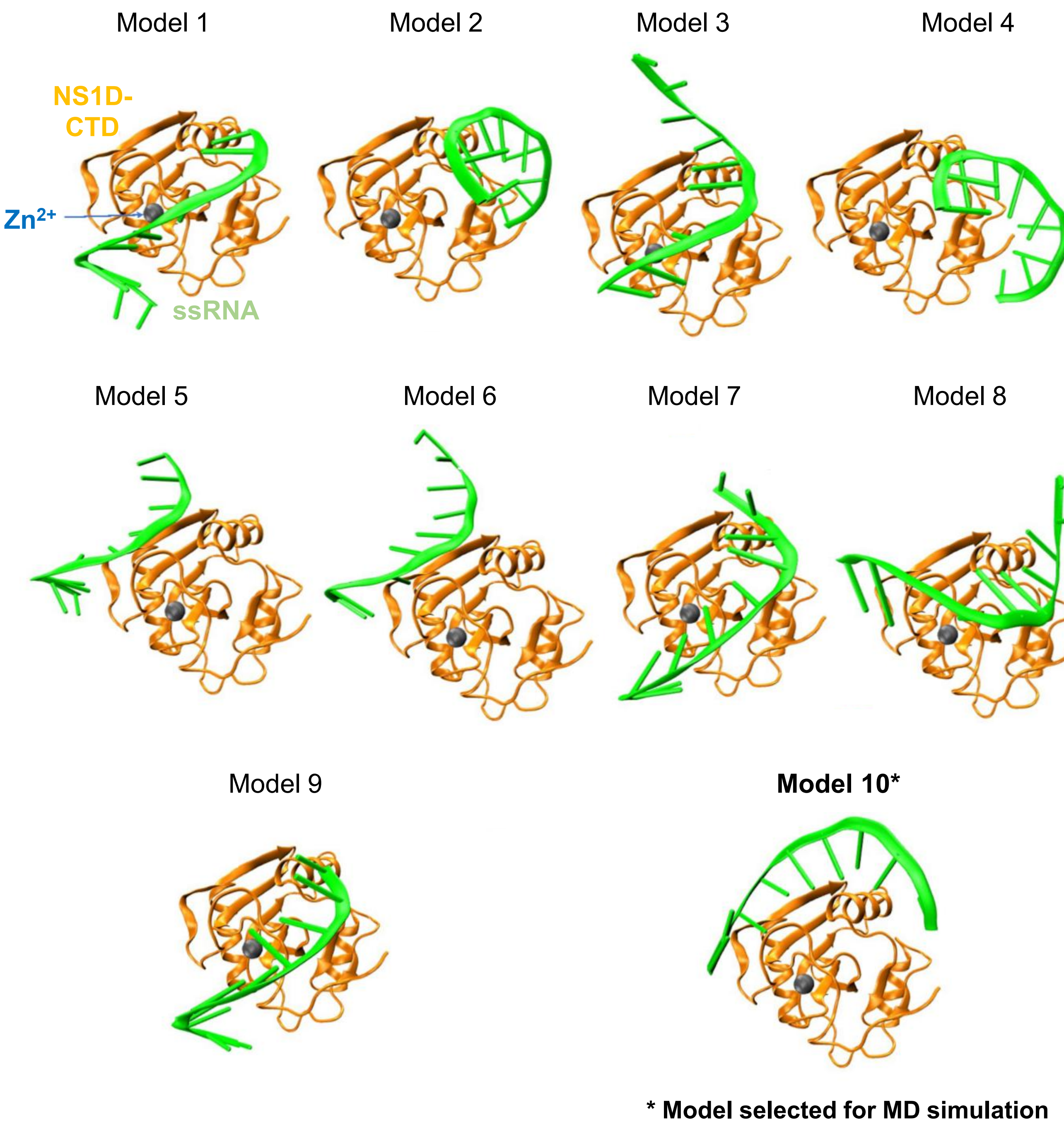


**S5 Fig. Ten highest-scoring docking models of the NS1D-CTD–ssRNA complex.** NS1D-CTD and ssRNA are shown as ribbon representations colored orange and green, respectively. Model 10 was selected for further molecular dynamics simulation because it most closely resembled the NS1D-CTD–ssRNA binding mode predicted by AlphaFold3.

(A)

| Oligo | Sequence | Alphafold 3 predicted scores | |
|---|---|---|---|
| | | ipTM | PTM |
| Scattered | CGUAUUCGUC | 0.72 | 0.86 |
| polyA | AAAAAAAAAA | 0.76 | 0.85 |
| polyU | UUUUUUUUUU | 0.81 | 0.88 |
| polyG | GGGGGGGGGG | 0.81 | 0.88 |
| polyC | CCCCCCCCCC | 0.7 | 0.83 |

(B)

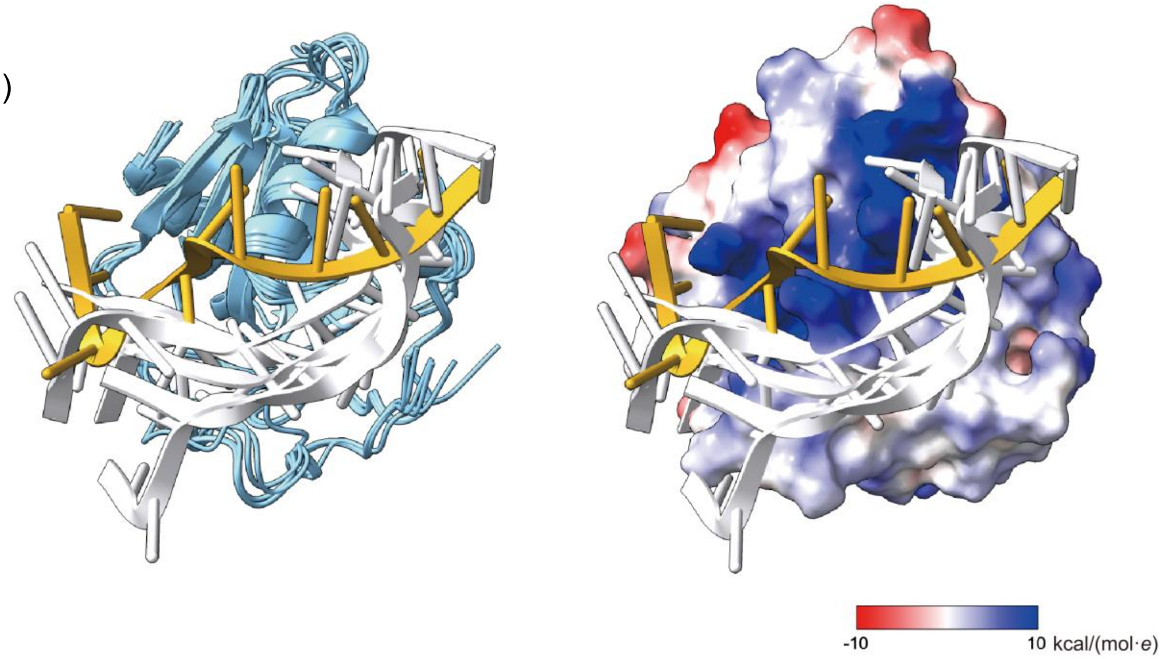


**S6 Fig. Structural modeling of NS1D-CTD-ssRNA complexes.** (A) Intrinsic model accuracy estimates (ipTM and pTM) for NS1D-CTD–ssRNA complexes predicted by AlphaFold-Multimer using five distinct ssRNA sequences. All predicted models achieved ipTM scores ≥0.70 and pTM scores >0.80, indicating consistently high confidence in the predicted protein–RNA interactions and overall complex structures. (B) Structural superimposition of the five AlphaFold3-predicted NS1D-CTD–ssRNA complexes with the HDOCK model selected for schematic representation of the NS1D-CTD–ssRNA interaction in this study. Despite differences in RNA sequence, all AlphaFold3 models and the selected HDOCK model after molecular dynamics simultation. The electrostatic surface potential of NS1D-CTD is shown with positively and negatively charged regions colored blue and red, respectively.

**S1 Table. Summary of the ten highest-scoring NS1D-CTD–ssRNA docking models.**

Summary of the Top 10 Models

| Rank | 1 | 2 | 3 | 4 | 5 | 6 | 7 | 8 | 9 | 10 |
|---|---|---|---|---|---|---|---|---|---|---|
| Docking score | -235.04 | -230.36 | -229.62 | -226.43 | -213.43 | -213.25 | -210.68 | -209.41 | -208.36 | -207.40 |
| Confidence score | 0.8456 | 0.8330 | 0.8310 | 0.8218 | 0.7819 | 0.7799 | 0.7709 | 0.7664 | 0.7626 | 0.7591 |
| Ligand rmsd (Å) | 32.57 | 30.40 | 23.54 | 31.49 | 39.24 | 34.34 | 34.80 | 23.15 | 38.29 | 31.44 |

“Note: Docking Score: The protein-protein/RNA/DNA complexes in the PDB normally have a docking score of around -200 or better. A more negative docking score means a more probable binding model. Confidence score: The docking score-dependent confidence score indicates the binding likeliness of two molecules. Roughly, when the confidence score is above 0.7, the two molecules would be very likely to bind; when the confidence score is between 0.5 and 0.7, the two molecules would be possible to bind; when the confidence score is below 0.5, the two molecules would be unlikely to bind”.
